# Reductive dehalogenation by diverse microbes is central to biogeochemical cycles in deep-sea cold seeps

**DOI:** 10.1101/2024.01.23.576788

**Authors:** Yingchun Han, Zhaochao Deng, Yongyi Peng, Jiaxue Peng, Lei Cao, Yangru Xu, Yi Yang, Hao Zhou, Chen Zhang, Dongdong Zhang, Minxiao Wang, Chunfang Zhang, Chris Greening, Xiyang Dong

## Abstract

Reductive dehalogenation is crucial for halogen cycling and environmental remediation, yet its ecological role is not completely understood, especially in deep-sea environments. To address this gap, we investigated the diversity of reductive dehalogenases (RDases) and ecophysiology of organohalide reducers in deep-sea cold seeps, which are environments rich in halogenated compounds. Through genome-resolved metagenomic analysis of 165 global cold seep sediment samples, we identified four types of RDases, namely prototypical respiratory, transmembrane respiratory, and cytosolic RDases, and one novel clade. Cold seeps were found to harbor a higher abundance of these RDase genes compared to other marine sediments, establishing them as unique hotspots for microbial reductive dehalogenation activity. These RDases are encoded by a wide range of microorganisms across four archaeal and 36 bacterial phyla, significantly expanding the known diversity of organohalide reducers. Halogen geochemistry, laboratory incubations with various halogenated compounds, metatranscriptomic data, and metabolomic profiling confirmed the presence of organohalides at concentrations of up to 18 mg/g in these sediments and demonstrated their active microbial reduction. This process is tightly linked to other biogeochemical cycles, including carbon, hydrogen, nitrogen, sulfur, and trace elements. RDases from cold seeps have diverse N-terminal structures across different gene groups, and reductive dehalogenase genes in these environments are mostly functionally constrained and conserved. These findings collectively suggest that reductive dehalogenation is a central process in deep-sea environments, mediated by a diverse array of microbes and novel enzymes.

## Introduction

Globally situated along continental margins, deep-sea cold seeps are regions where fluids rich in methane and other hydrocarbons migrate from the deep subsurface to sediment-seawater interface [1, 2]. These fluids support a diverse array of microorganisms, including anaerobic methanotrophic archaea (ANME) and their syntrophic sulfate-reducing bacterial partners [3, 4]. Beyond hydrocarbons, cold seeps are also rich in various organic compounds like organohalides. Various organohalides, such as dichloroacetaldehyde and tetrachlorobenzene, are abundant in cold seep sediments in various locations, including the Eastern Gulf of Mexico and the Haima cold seep in the South China Sea [1, 3, 5]. These organohalides, given their high standard redox potential (*Eo’* = +0.24 to +0.58 V), are desirable electron acceptors for anaerobic respiration in these oxidant-limited environments [1, 6, 7, 8, 9, 10, 11]. Organohalide-respiring microorganisms harness energy for growth by reducing these organohalides [6, 12, 13]. Notably some microorganisms within the *Chloroflexota* phylum, inhabiting deep-sea environments such as methane seeps, hydrothermal vents, and hadal trenches, can grow either obligately or facultatively on organohalides [14, 15, 16, 17]. However, the diversity and roles of organohalide-reducing microorganisms in deep-sea cold seeps remain largely uncharted.

The process of reductive dehalogenation is mediated by reductive dehalogenases (RDases, also termed as RdhA, encoded by *rdhA*), which catalyze the cleavage of halogen (like chloride or bromide) from the carbon backbone under anoxic conditions [18, 19]. For organohalide-respiring microorganisms, RDases are periplasmic membrane-associated proteins exported by the twin-arginine translocation (TAT) system [9, 19, 20]. These enzymes are typically composed of two subunits: the periplasmic catalytic subunit RdhA usually harbors a corrinoid cofactor that mediates halogen elimination and two tetranuclear [Fe-S] clusters for electron transfer, while the membrane-anchoring subunit RdhB typically contains two or three transmembrane helices (TMHs) [9, 19, 20]. The quinone-dependent RDases use respiratory quinols (e.g. menaquinol), reduced by electron donors such as molecular hydrogen (H2), whereas other enzymes are quinone-independent and appear to form respiratory supercomplexes with hydrogenases and potentially other primary dehydrogenases [19, 21, 22]. In addition to these prototypical RDases, two other classes of RDases are known. In the transmembrane respiratory RDases, the *rdhB* gene and TAT signal peptide are absent and instead the *rdhA* gene is directly connected to the membrane with one to three N-terminal TMHs [23, 24, 25]. In contrast, the cytosolic RDases (also called the catabolic RDases) are not connected to the respiratory chain and instead mediate the initial reductive dehalogenation of compounds so that they can be used as carbon and energy sources; these enzymes are typically NADPH-dependent and oxygen-tolerant, in contrast to their counterparts in organorespirers [19, 26]. However, classifying RDases remains challenging due to the limited number of well-characterized examples and their low sequence identity [9, 23, 26, 27, 28].

In this study, we analyzed a comprehensive dataset consisting of 165 metagenomic, 33 metatranscriptomic, and 55 metabolomic samples collected from deep-sea cold seeps globally. To evaluate the ecological significance of RDase resources and reductive dehalogenation in cold seeps, we also included public metagenomes from various marine sediment environments. Our investigation centered on the *rdhA* genes and organohalide reducers within cold seep ecosystems, systematically examining their phylogenetic diversity, ecological distribution, metabolic functions, genetic microdiversity, and structural evolution. To complement these analyses, we conducted laboratory incubations using sediment samples amended with various halogenated compounds to directly observe microbial dehalogenation activity.

We employed two major innovations in our study. First, given the limited number of experimentally solved RDase structures [18, 27, 29, 30], we employed AI-based protein structure modelling to better understand the structural and functional diversity within this superfamily. Second, we used state-of-the-art methods for quantifying genetic variations in microbial populations, including single-nucleotide variants (SNVs), to study the population dynamics of these enzymes across various cold seep environments [31, 32, 33, 34]. Through using a computational framework that integrates protein structure analysis with genetic variation [35], we were able to gain a more detailed understanding of the sequence-structure-function relationships of RDases, while uncovering a central role for organohalide reducers in shaping the biogeochemistry and ecology of deep-sea sediments.

## Results and Discussion

### Geochemical and experimental evidence for cold seep reductive dehalogenation

Reductive dehalogenation can cause an increase in halogen profiles in cold seep sediments [36]. Our analysis of 1,089 pore water samples (**Fig. 1A, Fig. S1 and Table S1**) revealed that the concentrations of dissolved chlorine (Cl⁻) and bromine (Br⁻) and the Br⁻/Cl⁻ ratio in cold seep pore waters are, on average, slightly higher than those in the overlying water and typical seawater [37, 38]. This trend is also observed in pore water iodide (I^-^) concentrations, which are higher in cold seep sediments compared to overlying seawater [39]. Additionally, the solid-phase concentrations of chlorine and bromine in 68 freeze-dried sediment samples showed a notable decrease with increasing sediment depth (**Fig. 1A and Table S2**), potentially suggesting reductive dehalogenation decreases with sediment depth. However, other geological and biological processes may also contribute to these differences, as indicated by the varied dissolved halogen profiles found in 84 cold seep sediment cores (**Figs. S2-S7)**.

**Fig. 1.**
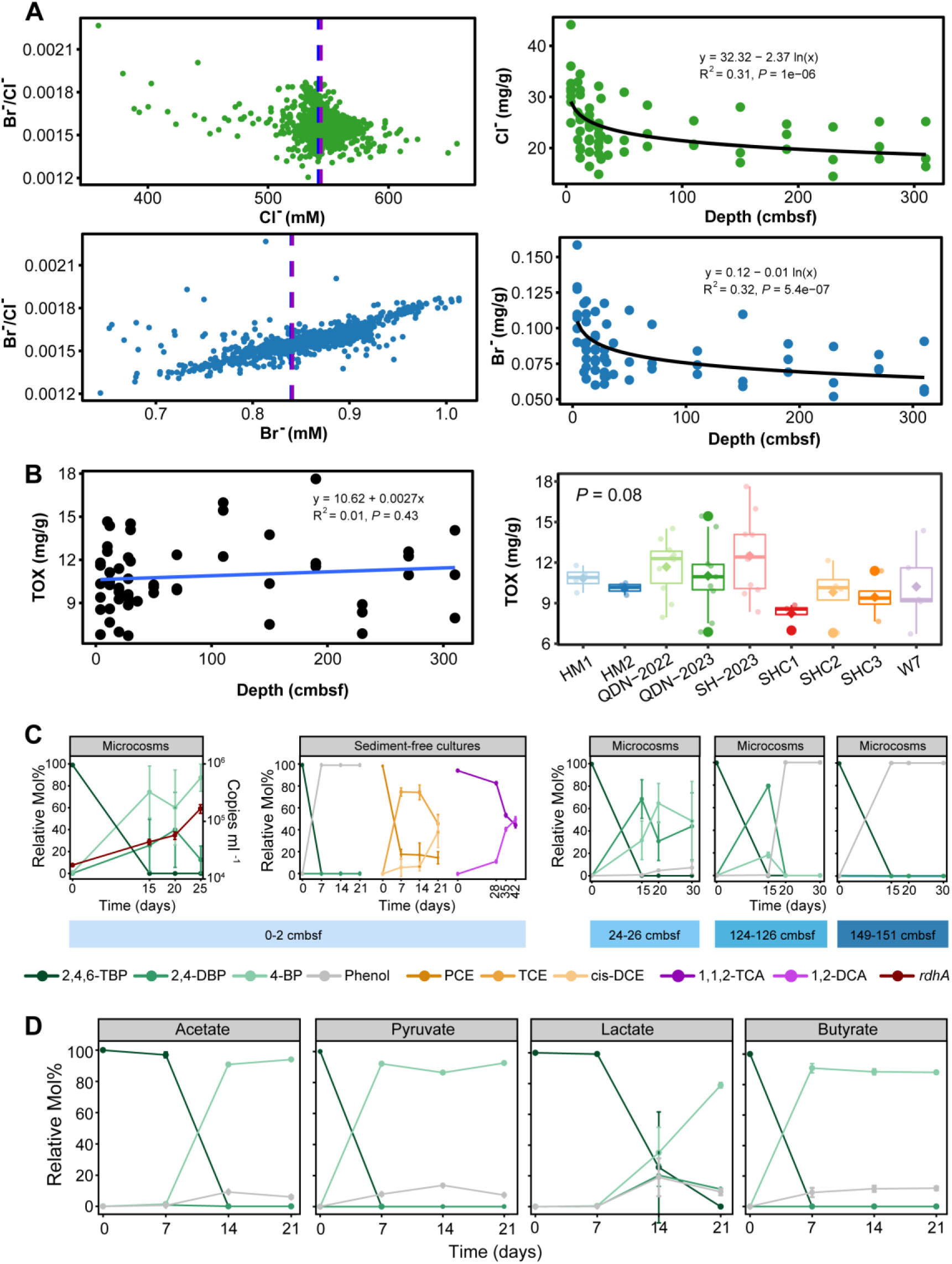
Halogen profiles in cold seep samples and degradation profiles of lab incubations. **(A)** Concentration profiles of dissolved (left) and solid (right) chlorine (Cl^-^) and bromine (Br^-^) in 1089 porewater samples (left) and 68 sediment samples (right) collected from Qiongdongnan, Shenhu, Haima, and Site F cold seeps. Blue lines and red lines represent the average concentration of Cl^-^ and Br^-^ in the typical seawater and overlying water of cold seeps, respectively. The depth of sediment samples is defined as centimeters below the sea floor (cmbsf). **(B)** Concentration profiles of total organic halogen (TOX) in sediments collected from Qiongdongnan and Shenhu cold seeps. The codes on the x-axis of the boxplot represent the different columns of cold seep sediments. **(C)** Degradation profiles of lab incubations. **Left:** Degradation profiles of 2,4,6-TBP and *rdhA* gene expression (black-red) in microcosms of surface sediments (0-2 cmbsf) over time, with relative mol% of 2,4,6-TBP (deep black-green), 2,4-DBP (green), 4-BP (light green), and phenol (gray) shown. **Middle:** Degradation profiles of PCE (yellow-orange) and 1,1,2-TCA (purple) in sediment-free cultures. **Right:** Degradation profiles of 2,4,6-TBP in microcosms at varying depths (24-26, 124-126, and 149-151 cmbsf) over time. Each microcosm or sediment-free culture was conducted in triplicate, with error bars representing the corresponding standard deviations. 2,4,6- TBP: 2,4,6-tribromophenol; 2,4-DBP: 2,4-dibromophenol; 4-BP: 4-bromophenol; TCE: trichloroethylene; cis-DCE: cis-1,2-Dichloroethylene; 1,2-DCA: 1,2-dichloroethane. **(D)** Debromination profiles of 2,4,6-TBP in sediment-free cultures with acetate, pyruvate, lactate, and butyrate as the sole electron donors.

In 55 cold seep sediment samples, total organic halogen (TOX) concentrations ranged from 6.7 mg/g to 17.6 mg/g, with an average of 10.9 mg/g. These concentrations showed no significant variations with sediment depth but differed among the columns of cold seep sediments (**Fig. 1B and Table S2**). Additionally, untargeted metabolomics analysis of these sediments identified 3,713 peaks, including 560 annotated metabolites. Among these metabolites (**Table S3**), a variety of halogenated compounds were detected, some reaching relative abundances of up to 10³based on peak intensity. These compounds included organochlorides (e.g., 2-chloro-L-phenylalanine), organobromides (e.g., bromobenzene-3,4-oxide), organofluorides (e.g., 5-fluorouridine triphosphate), and organoiodides (e.g., 5-iodo-2’-dUMP).

Incubation experiments using sediment samples from different depths confirmed dehalogenation activity in cold seeps **(Fig. 1C)**. In all four microcosms (0-2, 24-26, 124-126, and 149-151 centimeters below the sea floor [cmbsf]), 2,4,6-tribromophenol (2,4,6-TBP) was fully debrominated to 2,4-dibromophenol (2,4-DBP) and 4- bromophenol (4-BP) after 15 days (**Fig. 1C, left and right**). For the deeper samples (124-126 and 149-151 cmbsf), complete debromination to phenol was observed after 20 days. In parallel, sediment-free cultures also demonstrated the complete debromination of 2,4,6-TBP to phenol within 7 days (**Fig. 1C, middle**), suggesting that microbial debromination can occur without the involvement of electron mediators (e.g., humic substances) present in sediments [40, 41]. Further experiments showed that cold seep microorganisms could dechlorinate other halogenated compounds (**Fig. 1C, middle**), such as converting perchloroethylene (PCE) to trichloroethylene (TCE) and cis-dichloroethylene (cis-DCE), as well as transforming 1,1,2-trichloroethane (1,1,2- TCA) to 1,2-dichloroethane (1,2-DCA). No dehalogenation was observed in any of the abiotic controls, confirming the biological nature of these processes.

### Cold seeps harbor a vast repertoire of reductive dehalogenase genes with higher abundance than other marine sediments

Reductive dehalogenase (RdhA) sequences from cold seep metagenomes were identified and validated through analysis of a non-redundant gene catalog comprising 147 million genes [42], using a comprehensive workflow that incorporated sequence identity, gene length, conserved motifs, and phylogenetic validation (**Fig. S8**). The process yielded 3,993 validated RdhA sequences that fell into four phylogenetic groups **(Fig. 2**): (i) prototypical respiratory RDases (n = 1,466) associated with obligate and facultative organohalide-respiring microbes [19, 29, 43]; (ii) cytosolic RDases (n = 362) involved in liberating organohalides as energy and carbon sources [19, 26, 27]; (iii) transmembrane respiratory RDases (n = 1,963) characterized by a fusion of the *rdhA* gene with N-terminal TMHs [23, 26]; and (iv) a novel clade of RDases (n = 202). This clade, clustering between the cytosolic and prototypical clades, is characterized by the presence of both N-terminal TMHs (like transmembrane respiratory RDases) and a RdhB partner subunit (like prototypical respiratory RDases) **(Table S4)**. These four groups could be further subclassified into six subgroups, for example quinone-independent and quinone-dependent prototypical respiratory RDases, as depicted in **Fig. 2**.

**Fig. 2.**
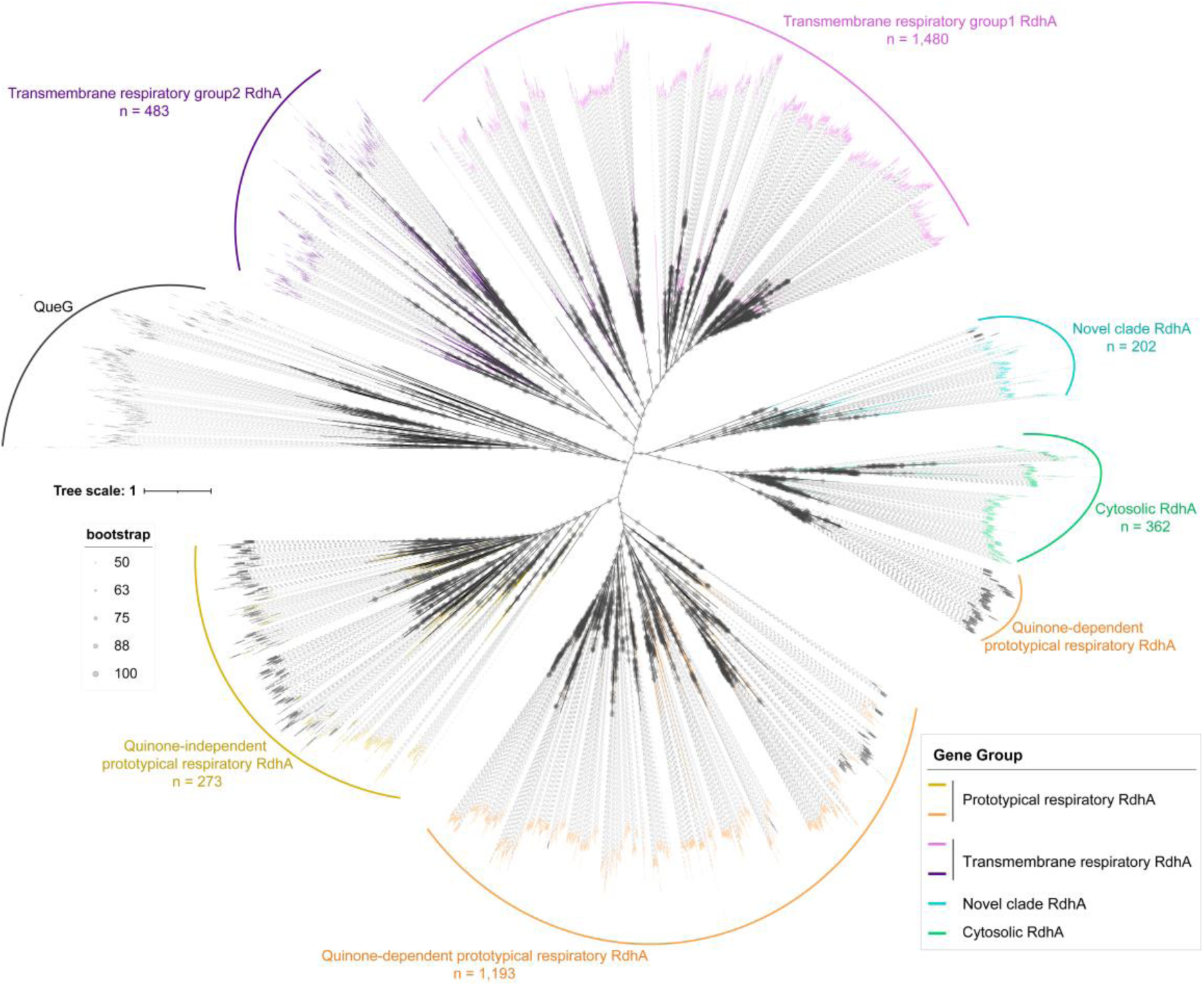
Maximum-likelihood phylogenetic tree of reductive dehalogenase (RdhA) protein sequences identified from the cold seep non-redundant gene catalog. Branches are color-coded to represent different RdhA subgroups. Epoxyqueuosine reductase (QueG) was used as the outgroup. Scale bar indicates the mean number of substitutions per site. Bootstrap values over 50% were shown next to the nodes. The n values denote the count of RdhA sequences within each subgroup.

The relative abundance of cold seep *rdhA* genes (averaging 25 genes per million [GPM]) was compared to those in other marine sediment environments, including the hadal zone (4 GPM), sinking particle (1 GPM), polar ocean (7 GPM), and marginal sea (7 GPM) (**Fig. S9**). Cold seeps exhibited the highest relative abundance of *rdhA* genes, suggesting that these environments serve as unique hotspots for microbial reductive dehalogenation activity. Additionally, cold seep *rdhA* genes showed statistically significant differences across the four groups and six subgroups (*P* < 2.2e-16; **Fig. 3A and Table S5**). Prototypical respiratory and transmembrane respiratory *rdhA* genes are more abundant than the cytosolic and novel clade *rdhA* genes (**Fig. 3A, inset**), suggesting that reductive dehalogenation in cold seeps is mostly associated with anaerobic respiration. Despite no notable differences in transcript abundance across groups (*P* > 0.05; **Fig. 3A**), the expression of *rdhA* subgroups (averaging 2.11 transcripts per million [TPM]; reaching 19.13 TPM for transmembrane respiratory *rdhA* genes) suggests microbial reductive dehalogenation occurs *in situ*. Furthermore, a prototypical respiratory *rdhA* gene was detected in the 2,4,6-TBP-dehalogenating culture, with its absolute expression level increasing tenfold (from 10^4^ to 10^5^ copies/ml) during the dehalogenation process (**Fig. 1C, left**).

**Fig. 3.**
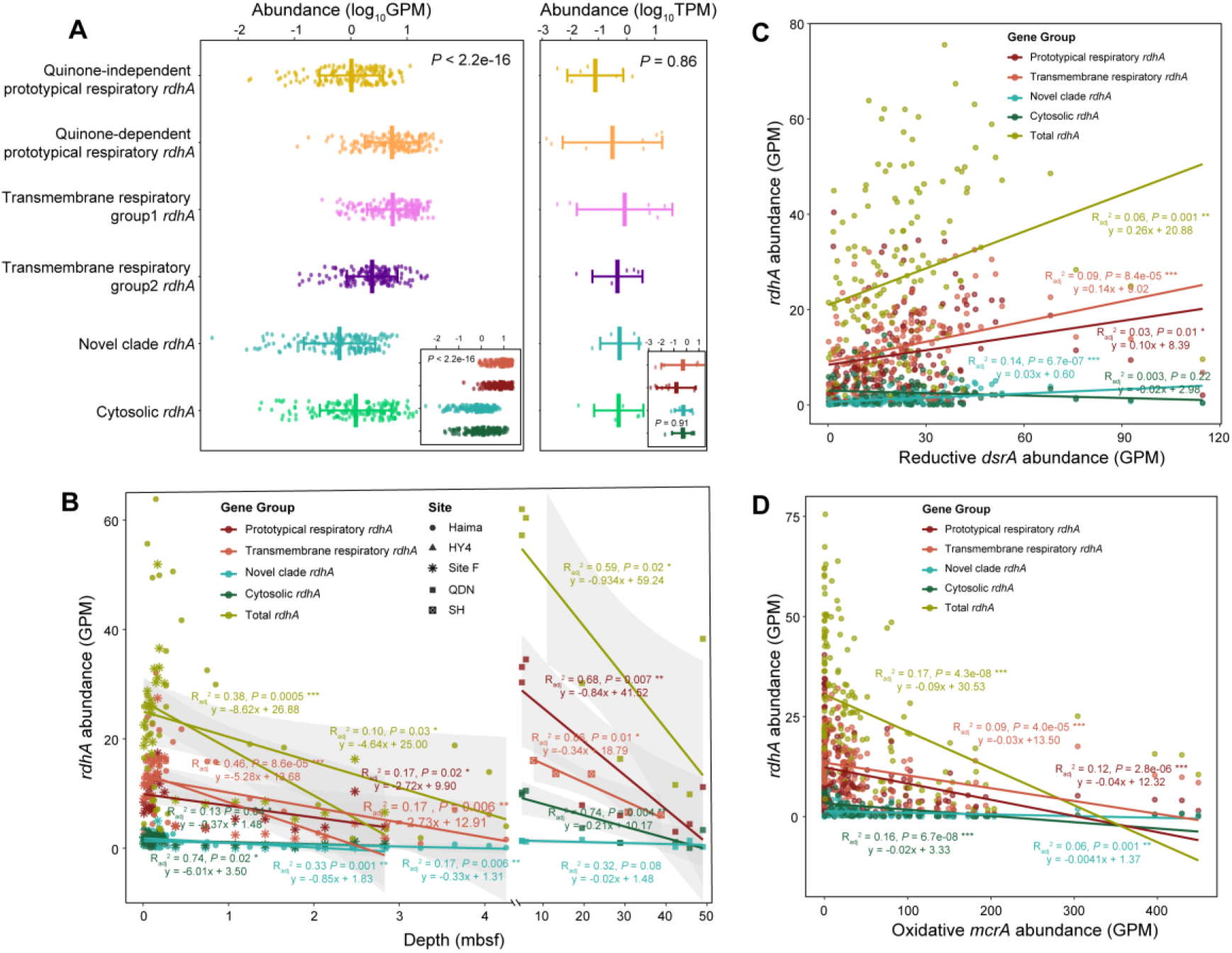
Relative abundance patterns of *rdhA* genes in cold seep sediments. **(A)** Relative abundance and expression levels of 3,993 *rdhA* genes across different sediment samples, measured as GPM (genes per million) for metagenomes and TPM (transcripts per million) for transcripts. Insert plots show the *rdhA* gene abundance and expression levels of the four groups: prototypical respiratory *rdhA* genes are shown in dark red, transmembrane respiratory *rdhA* genes in light red, novel clade *rdhA* genes in cyan, cytosolic *rdhA* genes in dark green. Gene groups and subgroups are in different colors. *P* values of differences across different types of *rdhA* genes were computed through Kruskal-Wallis rank-sum tests. Relationships between *rdhA* abundance and **(B)** sediment depths (mbsf), **(C)** reductive *dsrA* abundance, and **(D)** oxidative *mcrA* abundance. Each point represents the average gene abundance for a sample, accompanied by linear regression lines and *R*^2^ values specific to each gene group in corresponding colors. Detailed statistics for significance tests and linear regressions are provided in **Tables S5-S7**.

The distribution of *rdhA* genes appears to be influenced by sediment depth and cold seep types. A negative correlation between *rdhA* gene abundance and sediment depth (**Fig. 3B**) implies reduced dehalogenation activity in deeper sediments, potentially due to limited nutrients and bioavailable organohalides [44]. In contrast, some *rdhA* subgroups display site-specific, depth-dependent trends indicative of niche specialization (**Fig. S10A and Table S6**). Significant differences in *rdhA* gene abundance were also observed among five cold seep types, especially in quinone-independent prototypical respiratory, transmembrane respiratory and cytosolic *rdhA* genes (*P* < 0.05; **Fig. S11**).

Sulfate reduction and anaerobic methane oxidation, associated with *dsrA* and *mcrA* gene activities, respectively, are two important metabolic processes in cold seeps [45, 46]. The observed positive correlation between the abundances of *rdhA* and *dsrA* genes indicates that reductive dehalogenation and sulfate reduction processes likely occur simultaneously in cold seeps (**Fig. 3C and Table S7**). Moreover, the fact these genes are present in similar abundances suggests reductive dehalogenation is potentially as crucial as sulfate reduction in these environments, underscoring its significance in the biogeochemical dynamics of cold seeps. Conversely, the negative correlation with *mcrA* gene abundance (*P* < 0.001; **Fig. 3D and Fig. S12**) implies that anaerobic methane-oxidizing archaea may not engage in reductive dehalogenation [47, 48]. Additionally, the positive association of genes involved in osmotic stress protection and osmolyte transport with *rdhA* gene abundance (**Figs. S13-S14 and Table S8**) suggests that microorganisms deploy various salt tolerance mechanisms to maintain osmotic balance during reductive dehalogenation [49].

### Organohalide reducers span across four archaeal and 36 bacterial phyla

From the cold seep genome catalog with 3,164 MAGs (metagenome-assembled genomes), we identified 586 *rdhA* genes that each fell into the four groups (**Figs. S8 and S15**). These genes are distributed across 47 archaeal and 400 bacterial MAGs, encompassing four archaeal and 36 bacterial phyla, highlighting the wide distribution of organohalide reduction in these environments (**Fig. 4A, Fig. S16 and Table S9**). The most prevalent phyla harboring *rdhA* genes include *Chloroflexota* (n = 75), *Acidobacteriota* (n = 56), *Bacteroidota* (n = 55), *Desulfobacterota* (n = 53), *Krumholzibacteriota* (n = 28), and *Asgardarchaeota* (n = 27). Most MAGs (79%, n = 355) encoded a single *rdhA* gene, whereas 92 MAGs across 15 phyla contained multiple *rdhA* genes **(Fig. S16 and Table S9)**, in line with culture-based observations that highly versatile organohalide reducers produce multiple RDases [19, 29]. Further examination revealed 36 mobile genetic elements (MGEs) associated with 22 *rdhA*-containing contigs (**Table S10**), suggesting a role for MGEs in the horizontal transfer of *rdhA* genes and the resultant diversity of organohalide reducers. Additionally, the abundance of these organisms negatively correlates with sediment depth, in line with the decrease in *rdhA* gene abundance in deeper sediment layers (**Fig. 3B and Fig. S17**). To investigate potential substrates for organohalide reducers, we conducted molecular docking for RdhAs against 191 varied naturally occurring organohalides, including those commonly found in cold seeps **(Table S11)** [3, 5, 14]. In total, we found 13,561 possible interactions between RdhAs and halohydrocarbons, with the binding energies ranging from -0.47 to -6.08 kcal/mol **(Table S12)**. These interactions were further validated by the successful anaerobic microbial dehalogenation of aromatic and aliphatic compounds such as 2,4,6-TBP, 2,4-DBP, 4-BP, TCA, TCE, and PCE in sediment-free cultures (**Fig. 1C, middle**).

**Fig. 4.**
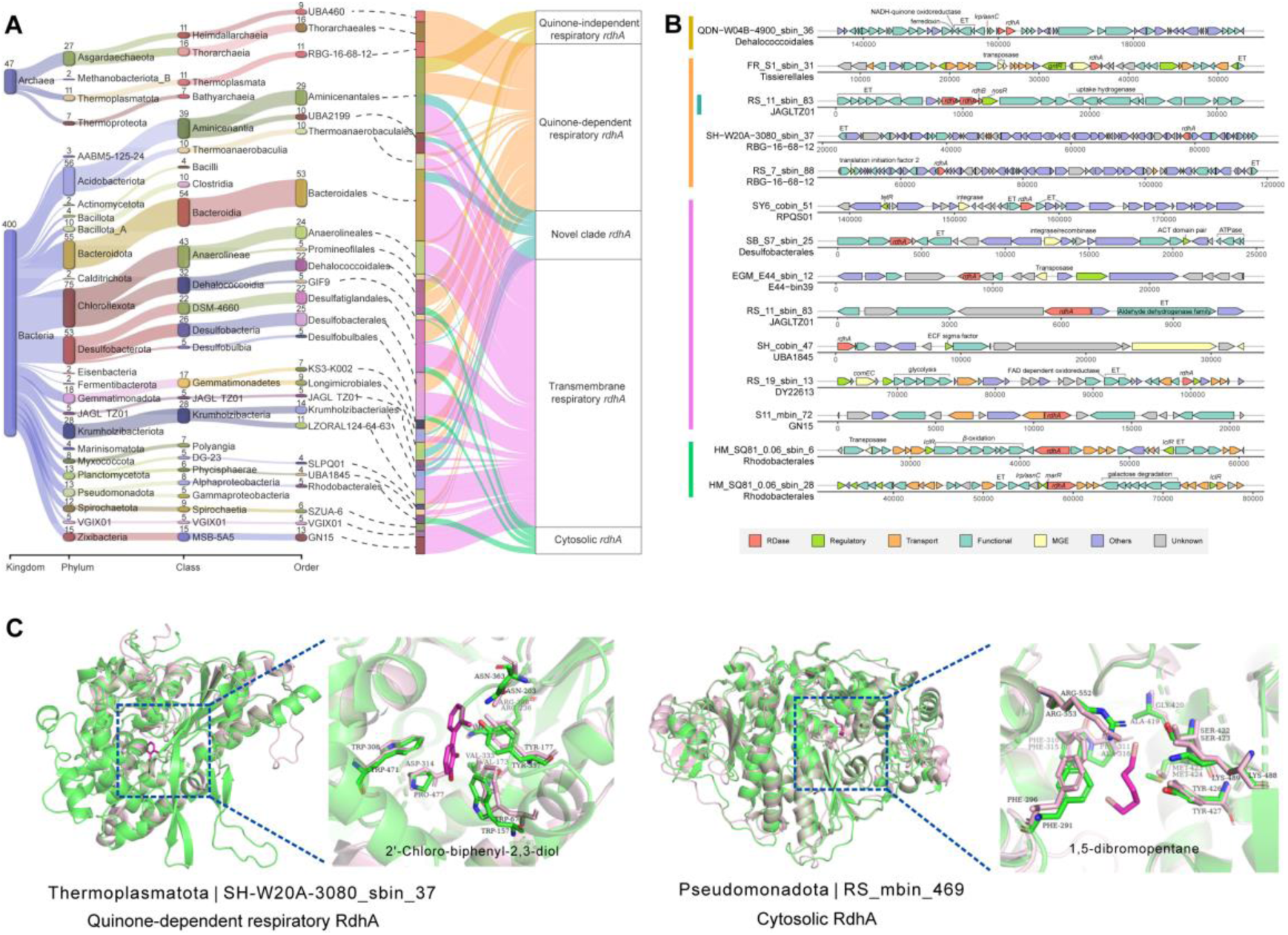
Taxonomic classification, genomic context and potential substrate docking of cold seep organohalide-reducing microorganisms. **(A)** A Sankey diagram showing the taxonomy of archaeal and bacterial MAGs according to GTDB, categorizing organohalide-reducers across taxonomic levels. **(B)** Genome synteny of 14 contigs showcasing five types of RDases (highlighted in red) and accompanying genes: regulatory (green), transport-related (orange), functional (blue), mobile genetic elements (MGEs, yellow), other annotated genes (purple), and unknown (grey). **(C)** Overlay of the active sites of cold seep quinone-dependent respiratory and cytosolic RdhA (lightpink) against known RdhA crystal structures (PDB id: 8Q4H and PDB id: 6ZY1, in green). Substrates 2’-Chloro-biphenyl-2,3-diol and 1,5-dibromopentane, predicted through molecular docking with binding affinities of -4.12 and -3.11 kcal/mol respectively, are shown in magenta. Details for taxonomic classification, molecular docking, and *rdhA*-containing contig annotations are provided in **Tables S9 and S11- S13**.

The quinone-dependent respiratory *rdhA* genes were encoded by 136 microorganisms spanning 15 bacterial and four archaeal phyla, including most of the putative archaeal organohalide reducers (**Fig. 4A, Fig. S16 and Table S9**). These RdhAs show strong structural homology to the PceA dehalogenase (PDB id: 4UR3) from *Sulfurospirillum multivorans* [18], with TM-scores (Template Modeling scores) between 0.63 and 0.90 (**Fig. S18**). In addition to be co-encoded with the membrane anchor subunit RdhB, this RDase is also frequently genomically associated with a partner uptake hydrogenase (Hup), the ferredoxin:NAD^+^ oxidoreductase (Fnr), and transcriptional regulators (for example, GntR family, and NosR/NirI family) (**Fig. 4B and Table S13**). Molecular docking analysis against 191 organohalides, including those abundant in cold seeps, demonstrated that *Thermoplasmata* RDase has a binding affinity of -4.1 kcal/mol for 2’-chloro-biphenyl-2,3-diol and thus is a possible substrate (**Fig. 4C and Table S12**). Quinone-independent respiratory *rdhA* genes, typically found in obligate organohalide reducers *Dehalococcoides* and *Dehalogenimonas* within the phylum *Chloroflexota* [6, 19, 29], were also detected in *Acidobacteriota* and *Gemmatimonadota* **(Fig. 4A and Fig. S16)**, broadening the known diversity of microorganisms harboring this gene type. Adjacent genes to *rdhA*, ferredoxin and NADH-coupled oxidoreductase were identified **(Table S13)**, which are essential for the electron transport chain in organohalide respiration [50]. RDases in *Aminicenantaceae* (*Acidobacteriota*) and GWA2-58-10 (*Gemmatimonadota*) exhibit -4.0 to -4.8 kcal/mol binding affinities for chlorinated benzene and chlorinated paraffins **(Table S12)**, suggesting their suitability as substrates.

The transmembrane respiratory RDases were by far the most widespread enzymes, encoded by 287 microorganisms from 33 phyla, including multiple *Bacteroidota*, *Acidobacteriota*, *Desulfobacterota* and *Krumholzibacteriota* MAGs. This observation extends Atashgahi et al.’s findings that these enzymes are widespread despite being understudied [23]. Adjacent genes to *rdhA*, transcriptional regulators (e.g., GntR family), genes involved in electron transport (e.g., FAD dependent oxidoreductase) and MGEs (e.g., transposase) were identified **(Fig. 4B and Table S13)**. We also observed transmembrane respiratory RDases form various phyla exhibit -3.2 to -6.0 kcal/mol binding affinities for chlorinated benzene and chlorinated alkane **(Table S12)**, suggesting potential substrates.

The novel clade was also widespread, encoded by 49 MAGs spanning 11 phyla, including the four abovementioned phyla and candidate phyla JAGLTZ01 and QNDG01 (**Fig. 4A, Fig. S16 and Table S9**). Despite being phylogenetically distant from the transmembrane respiratory RDases, this novel clade also likely has a respiratory function, given the *rdhA* gene is typically fused with between one to three N-terminal TMHs; however, a key difference is that 23 and 38 of the 50 genes in the novel clade are sometimes found alongside TAT systems and *rdhB* genes, respectively **(Table S9)**. Novel RDases demonstrate binding affinities ranging from -4.4 to -6.1 kcal/mol for chlorinated benzene **(Table S12)**, indicating their suitability as substrates.

Cytosolic *rdhA* genes (n = 37) were mainly found in *Chloroflexota*, *Desulfobacterota*, and *Pseudomonadota*, but were also detected in a *Heimdallarchaeia* MAG **(Fig. 4A, Fig. S16 and Table S9)**. Consistent with a role in organic carbon degradation, these genes are often associated with bacterial transcriptional regulators IclR/MarR, beta oxidation genes, and galactose degradation genes **(Fig. 4B)**. In three MAGs, cytosolic *rdhA* genes were discovered on plasmids (**Table S14**); this is reminiscent of BhbA, the plasmid-encoded first characterized enzyme from this family, which aerobically breaks down bromoxynil into 4-carboxy-2-hydroxymuconate-6-semialdehyde in *Comamonas* sp. 7D-2, which is then funneled into the tricarboxylic acid cycle [19, 51]. These cytosolic RdhAs have high TM-scores (0.92 to 0.95) (**Fig. S19**) to the X-ray crystal structure of the 3-bromo-4-hydroxybenzoic acid reductase from *Nitratireductor pacificus* (NpRdhA; PDB id: 6ZY1) [27]. Two organohalide reducers from *Gammaproteobacteria*, encoding cytosolic RDases, showed potential in reducing 1,5- dibromopentane, evidenced by binding energies of -2.44 and -3.11 kcal/mol (**Table S12**). Further analysis revealed that these microorganisms contain genes responsible for the aerobic degradation of medium-chain alkanes (C5-C13, e.g. pentane; *alkB* and CYP153) **(Fig. 5 hydrocarbon degradation and Table S15)**, suggesting a link between catabolic reductive dehalogenation and aerobic hydrocarbon degradation [19, 27, 51].

**Fig. 5.**
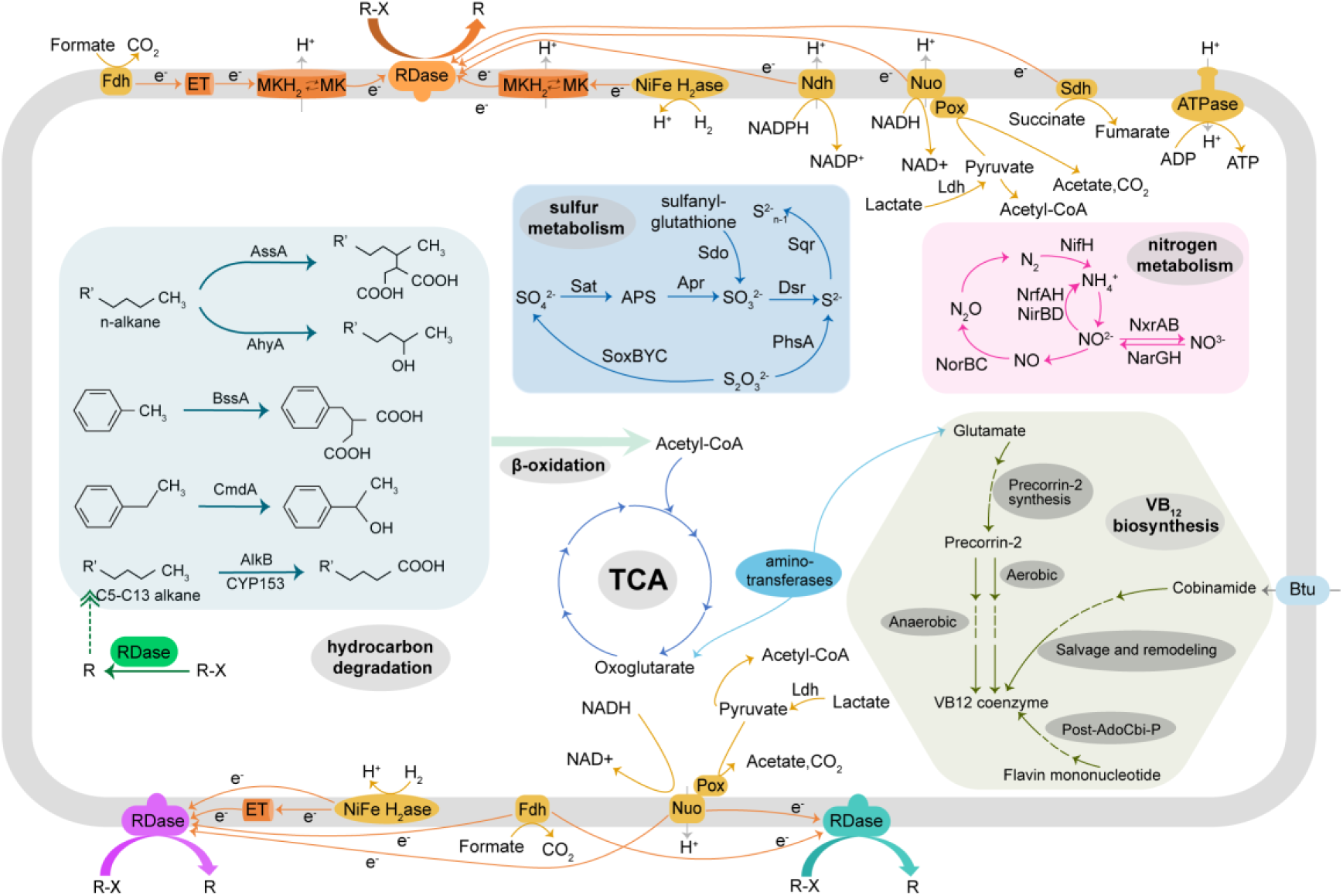
Reductive dehalogenation is closely linked to carbon, nitrogen, sulfur, and trace element metabolic processes. RDases of different colors represent different types, with orange indicating prototypical respiratory RDases, purple representing transmembrane respiratory RDases, green indicating cytosolic RDases, and cyan signifying the novel clade of RDases. Dark orange denotes potential electron transfer units, while orange-yellow represents potential electron donors and electron transfer pathways. Different background colors signify distinct metabolic pathways: light blue indicates hydrocarbon degradation, dark blue represents sulfur metabolism, light red represents nitrogen metabolism and olive green denotes B12 biosynthesis. Detailed annotations for different genes are provided in **Tables S14-S19.**

### Cold seep halogen cycling is closely linked to central biogeochemical processes

We extensively annotated the metabolic capabilities of the RDase-encoding MAGs to gain insights into their potential ecophysiology and biogeochemical roles. Fermentation is the main process transforming organic carbon in cold seep environments [1, 52], which produces significant amounts of hydrogen (H2) and various organic acids, both of which are key electron donors for organohalide reducers [43, 49, 53, 54]. Consistently, 288 organohalide reducers were found to encode 495 hydrogenases. Of these, 337 are involved in H2 oxidation [55], including group 1a, 1b, 1c, 1d, 1e, and 1f [NiFe]-hydrogenases that donate H2-derived electrons to anaerobic respiratory chains **(Fig. S20 and Table S16)**. Additionally, genes responsible for formate (*fdhAB*, *fdoG*, *fdwB* and *fdoH*), acetate (*acs*, *acdA*, *ack* and *pta*), pyruvate (*pdh*, *aceE*, *ofor*, *por*, *poxB* and *poxL*), and lactate (*ldh*) oxidation were present in 226, 363, 448, and 18 organohalide reducers, respectively (**Fig. 5, Fig. S20 and Table S17**). We observed that different electron donors (acetate, pyruvate, lactate, and butyrate) affected the debromination process of 2,4,6-TBP in cold seep enriched cultures (**Fig. 1D**). Notably, during the first 7 days, the average bromide removal rate was approximately 2.5 μM d^-1^ in the pyruvate and butyrate groups but decreased dramatically to less than 0.1 μM d^-1^ in the acetate and lactate groups. Despite this substrate specificity, the sustained debromination activity across various electron donors underscores the link between reductive dehalogenation and organic acid metabolism. Most of organohalide reducers involved in processing hydrogen and organic acids possess RdhAs that belong to prototypical and transmembrane respiratory RDases.

Organohalide reducers also participate in nitrogen, sulfur, and hydrocarbon cycles **(Fig. 5 nitrogen and sulfur metabolism)**. We identified 55 and 57 organohalide reducers with genes related to sulfate and thiosulfate reduction (reductive *dsrA* and *phsA*), respectively (**Tables S17-S18**). Additionally, 185 organohalide reducers harbor genes involved in the oxidation of sulfide, sulfur, and thiosulfate (*sqr*, *sdo* and *soxBYC*; **Table S17**). 189 organohalide reducers harbor reductive genes associated with nitrogen metabolism (*narGH*, *nrfHA*, *nirBDKS* and *octR*), with a small proportion also capable of nitrogen fixation (n = 6) and nitrite oxidation (n = 5). Four organohalide reducers also encoded enzymes for the anaerobic degradation of hydrocarbons (**Fig. 5 hydrocarbon degradation, Fig. S20 and Table S15)**. Two of these, affiliated with *Desulfobacterota* and encoding quinone-dependent respiratory RDases, encode genes for the anaerobic degradation of n-alkanes (*assA*) [3, 56, 57]; they potentially remove chlorine from chlorinated paraffins, as suggested by their binding energy of -4.4 kcal/mol **(Table S12)**. Altogether, the presence of these genes indicates that organohalide reducers are not only versatile in their metabolic functions but also well-equipped to adapt and thrive in various environmental conditions. Despite previous findings suggesting the coupling of anaerobic oxidation of methane (AOM) with reductive dehalogenation [7], *rdhA* genes were not found in the ANME group of microorganisms **(Fig. S21 and Table S9)**. Moreover, although 55 sulfate-reducing bacteria have the capacity for reductive dehalogenation, none seem to partner with ANME in this process (**Table S18**). Considering the negative correlation between the gene abundance of *rdhA* and oxidative *mcrA* **(Fig. 3D)**, we hypothesize that these two processes are likely not coupled within cold seeps.

Cobalamin (vitamin B12) commonly acts as a cofactor for RDases, facilitating electron transfer [27, 29, 58]. Only 19% (n = 87) of organohalide reducers are capable of B12 biosynthesis through various pathways [59], including the aerobic (n = 2), anaerobic (n = 5), salvage remodeling (n = 12) and the Post-AdoCbi-P (n = 78) pathways **(Fig. 5, Fig. S20 and Table S19**). This finding is in line with existing knowledge that only a limited number of microbial genera in the ocean can synthesize B12 de novo [60, 61]. Among two capable of aerobic B12 biosynthesis, they encode cytosolic *rdhA* genes. The remaining 80% (n = 360) of organohalide reducers, which lack B12 biosynthesis genes, likely rely on B12 synthesized by other microorganisms or present in the environment for their reductive dehalogenation activities [62].

### Cold seep RDases show diverse N-terminal structures across different gene groups

Consistent with the phylogenetic tree based on protein sequences, the structure-based phylogeny also supports the classification of cold seep RDases into four distinct groups (**Fig. 6 and Table S20**): prototypical respiratory RDases, transmembrane respiratory RDases, cytosolic RDases, and the novel clade RDases. Structurally, bacterial and archaeal RDases from all four groups each contain corrinoid- and FeS-containing domains, each marked by a two-layered alpha-beta structure [18, 27, 29, 30]; however, these enzymes greatly vary in their N-terminal structures in a manner consistent with their physiological roles. Our structural phylogeny revealed key aspects of RDase evolution that sequence analysis alone could not, suggesting at least five key evolutionary events (**Fig. 6**). For the novel clade, the emergence of a single N-terminal α-helix separated this group from other RDases (**Event 1 in Fig. 6**). Most novel clade RDases (36 of 50) contain a single N-terminal transmembrane α-helix (**Fig. S22A**), likely anchoring these enzymes for a respiratory function [63]. Half of these novel clade RDases (18 of 36) with a single TMH also possess RdhB subunit, and the combination of the two forms a multi-transmembrane helix predicted by AlphaFold Multimer (**Fig. S23**). Other RDases within this novel clade anchor to the membrane through the RdhB subunit (**Fig. S22B**).

**Fig. 6.**
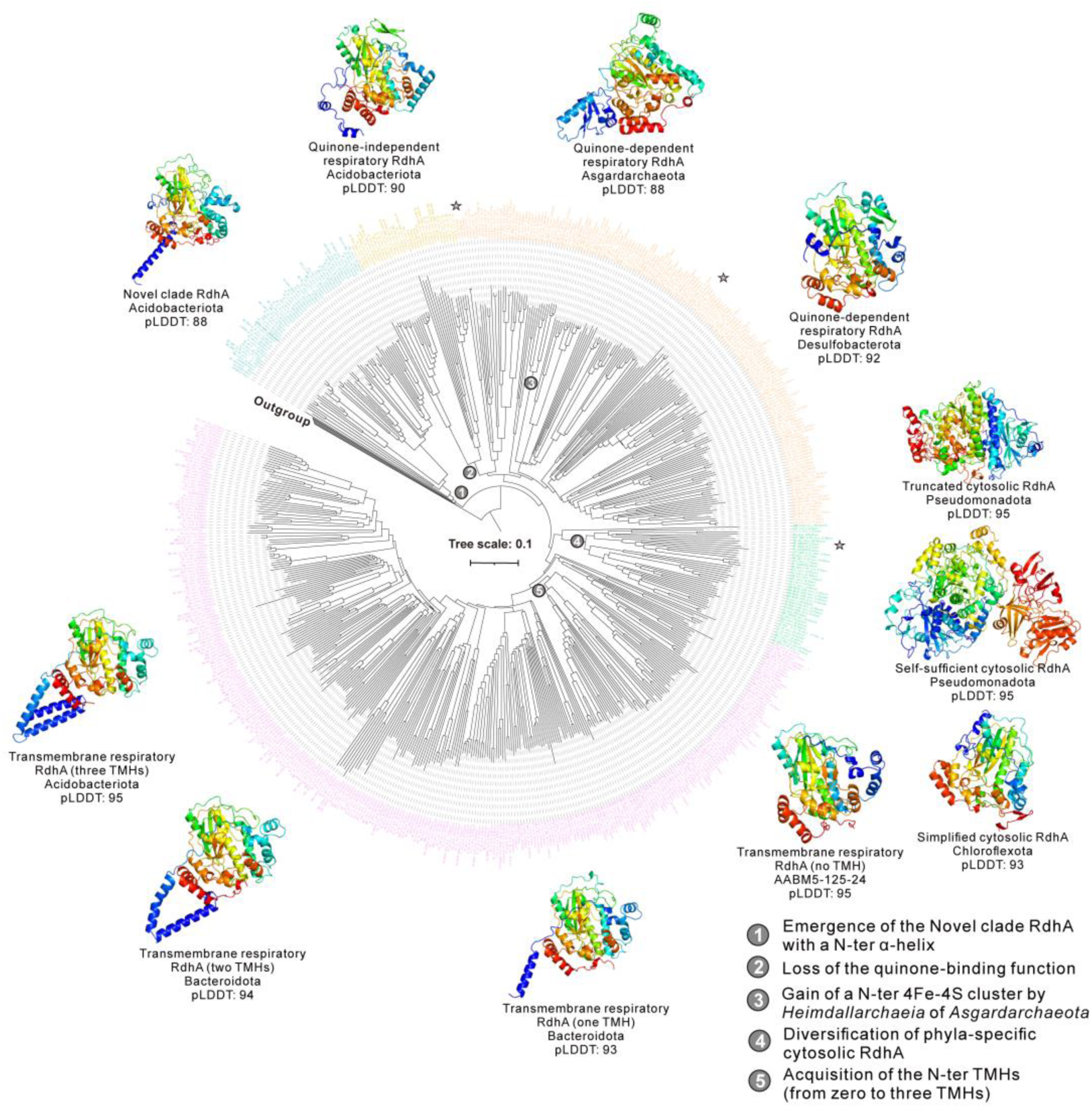
Structure-based phylogeny of cold seep reductive dehalogenases and reference proteins. Each group of reductive dehalogenases (RDases) is labeled by a unique color: Novel clade RDases, turquoise; Quinone-independent respiratory RDases, gold; Quinone-dependent respiratory RDases, orange; Cytosolic RDases, green; Transmembrane respiratory RdhAs, purple. The three RDases that have been validated experimentally are highlighted with stars. Surrounding the tree, the AlphaFold-predicted 3D structures of representative RDases are displayed, each labeled with their taxonomic phylum and corresponding pLDDT scores. The indication of the five main inferred evolutionary events of the tree marked at the origin of each fold/architecture, numbered 1 through 5.

Of the prototypical respiratory RDases, both the quinone-dependent and quinone-independent enzymes share structural similarities. They each contain a domain analogous to the quinone-binding domain found in quinone-dependent enzymes **(Fig. 6)**, such as the DhPceA2B2 from the tetrachloroethene-dechlorinating strain *Desulfitobacterium hafniense* TCE1 [29]. However, this domain in quinone-independent RDases does not exhibit quinone-binding affinity, in contrast to their quinone-dependent counterparts which show a significant binding affinity (-6.39 kcal/mol) and possess shorter substrate transport tunnels (**Event 2 in Fig. 6 and Fig. S24**). Additionally, among the eleven RDases from *Heimdallarchaeia*, five quinone-dependent RDases have complete N-terminal 4Fe-4S ferredoxin domains, with three others having incomplete ones (**Event 3 in Fig. 6, Fig. S25 and Table S9**). The presence of an additional 4Fe-4S domain in these archaeal RDases could potentially enhance the efficiency of the dehalogenation process by using multiple electron pathways from the quinone pool to the active sites [8, 29].

Cytosolic RDases are variable in length, ranging from 300 to 1000 amino acids, and they cluster into three distinct groups correlated with their respective phyla (**Event 4 in Fig. 6 and Table S9**). Eight cold seep cytosolic RDases, predominantly found in the *Pseudomonadota* phylum (with more than 700 amino acids), function as monomers due to the presence of an additional vestigial cobalamin-binding core domain [26]. Half of these cytosolic RDases are characterized by a unique C-terminal NADP-linked ferredoxin reductase domain [26], with an alpha-beta three-layered sandwich structure, allowing electrons derived from NAD(P)H to drive reductive dehalogenation (**Table S20**). According to the presence or absence of C-terminal reductase domain, these cytosolic RDases are termed as self-sufficient and truncated cytosolic RDases [26]. Another clade of cytosolic RDases (labelled as simplified cytosolic RDases) is typically present in *Desulfobacterota* and *Chloroflexota*, with an average length of 395 amino acids (**Fig. S26**). Simplified cytosolic RDases possess a shortened N terminus compared to both self-sufficient and truncated cytosolic RDases, as they lack the vestigial cobalamin-binding domain, similar to prototypical respiratory RDases. The considerable variation in the length of cytosolic RdhA proteins indicates significant genetic diversity, which may be indicative of their functional adaptability among various microbial taxa.

In the transmembrane respiratory RdhA proteins, the presence of an integral membrane domain is a common feature (**Event 5 in Fig. 6**), yet not all members of this group exhibit N-terminal TMHs. Specifically, certain RdhAs in the two earliest clades within this group lack transmembrane α-helices (**Fig. 6**). Nevertheless, approximately half of these proteins compensate for this absence by utilizing transmembrane domains located in adjacent genes for membrane anchoring (**Table S9**). The number of α-helices within these transmembrane domains varies, with some having one to three α-helices (**Fig. S27**), which likely impacts how these proteins embed in the membrane and potentially in turn their substrate specificity and catalytic efficiency.

### Cold seep *rdhA* genes are mostly functionally constrained and conserved

To investigate the evolutionary ecology of *rdhA* genes in cold seeps, we analyzed their genetic diversity across various sediment sites. Nucleotide diversity (within-population genetic diversity) across *rdhA* genes was consistently low, with an average of 0.013 (**Fig. 7A and Table S21**). Most *rdhA* genes were under strong purifying selection, as evidenced by low pN/pS values (the ratio of nonsynonymous to synonymous mutations; average 0.21 with 91.4% of ratios less than 0.4), suggesting reductive dehalogenation provides an adaptive advantage *in situ*. However, some quinone-dependent prototypical and cytosolic *rdhA* genes showed evidence of positive selection (**Table S21**), potentially due to genetic drift or beneficial mutations [64]. Further 3D structural predictions for *rdhA* genes with high pN/pS values (>1.5) [65] demonstrated that amino acid changes related to non-synonymous mutations did not significantly alter overall structure or catalytic domains of the proteins (**Fig. S28**). Additionally, the variation in nucleotide diversity and pN/pS ratios was statistically significant across different *rdhA* types (*P* = 2.2e-13 and *P* < 2.2e-16, respectively; **Fig. 7A and Table S21**), indicating unique evolutionary trajectories for each *rdhA* group. Nucleotide diversity also varied significantly among different types of cold seeps (*P* = 0.0006), with gas hydrates, oil and gas seeps, and methane seep samples exhibiting higher nucleotide diversity (**Fig. S29A**). Nevertheless, the pN/pS ratios were consistent across the five cold seep types (*P* = 0.25; **Fig. S29B**). Overall, these findings are in line with previous observations of key functional genes in microbial populations within cold seeps [31, 66], highlighting the strong functional constraints and conservation of essential metabolic *rdhA* genes across various cold seep environments [67].

**Fig. 7.**
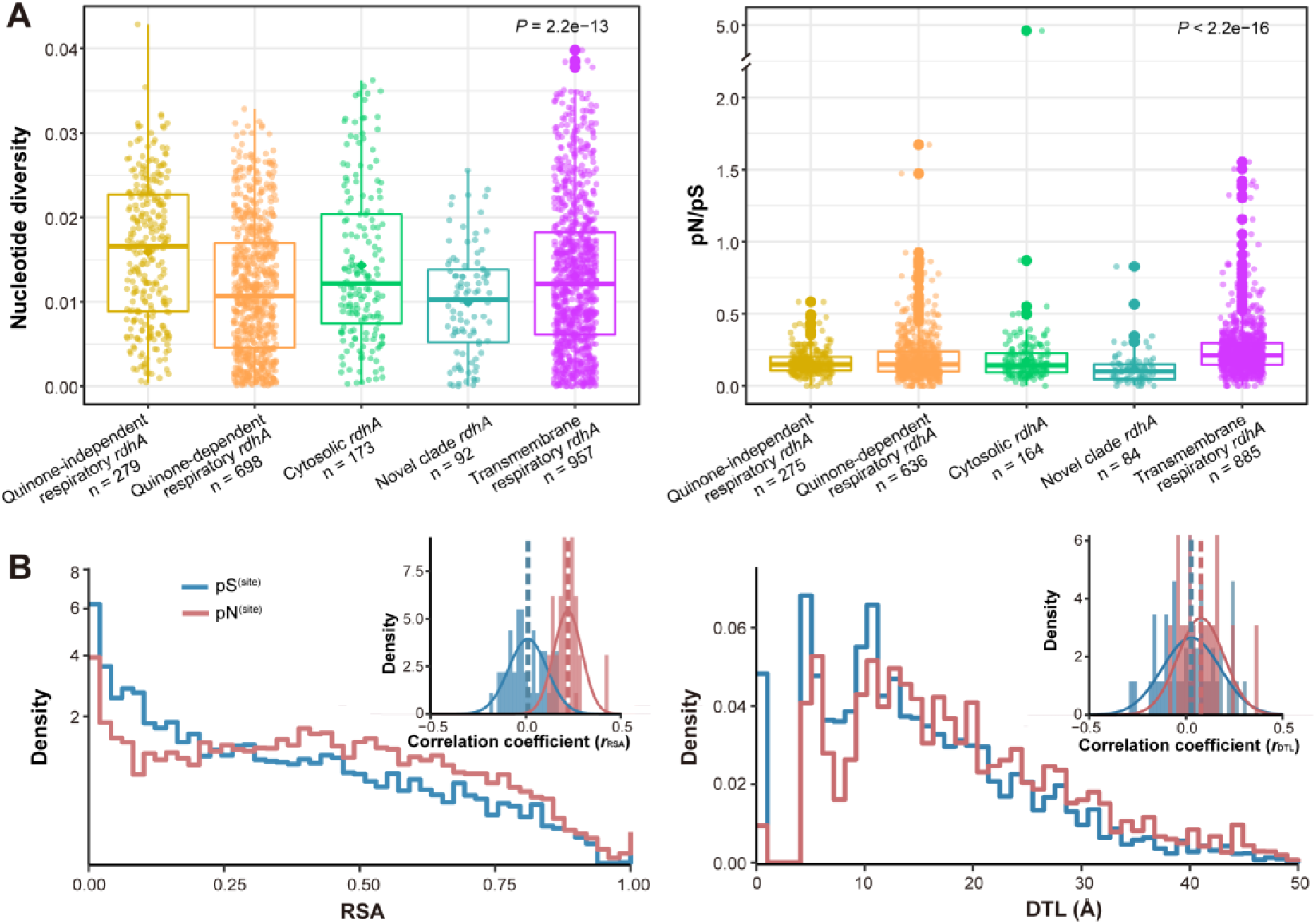
Microdiversity of cold seep *rdhA* genes and their corresponding protein structural features. **(A)** Comparison of nucleotide diversity and pN/pS for *rdhA* genes across gene groups. **(B)** Gene-wide distributions and Pearson correlations for pN^(site)^ (red) and pS^(site)^ (blue) relative to relative solvent accessibility (RSA) and distance to ligand (DTL). The pN^(site)^ and pS^(site)^ distribution were generated based on RSA and DTL values of each site from 200 predicted RdhA structures. The Pearson correlations of log10(pN^(site)^) and log10(pS^(site)^) to RSA and DTL for each gene-sample pair were calculated using linear models, and the mean values were represented as dashed lines. *P* values of differences across different types of *rdhA* genes were computed through Kruskal-Wallis rank-sum tests. The boxplot features include: center lines, medians; box limits, 25th and 75th percentiles; whiskers, 1.5× interquartile range from the 25th and 75th percentiles; points, outliers. The n values denote the number of independent results utilized for statistical derivation.

To further elucidate the association of nucleotide polymorphisms and protein structures of RDases in cold seeps, genetic variation and biophysical characteristics of RDases were examined. This included single-codon variant (SCV), synonymous (s) and nonsynonymous (ns) polymorphism rates of each codon (pS^(site)^ and pN^(site)^), as well as evaluating relative solvent accessibility (RSA) to indicate the exposure or burial of a site, and distance to ligand (DTL) to represent the proximity to the nearest active site at each codon position, as recently described in Kiefl et al. [35]. A total of 50,531 SCVs (1,684 per metagenome on average) were found, and pS^(site)^ exceeded pN^(site)^ by a ratio of 6:1 on average of all SCVs, exhibiting particularly high enrichment of synonymous polymorphism within the *rdhA* genes. Additionally, pN^(site)^ and pS^(site)^ values varied significantly from site to site (78.33% ± 0.13% versus 79.75% ± 0.11% of total variance, ANOVA), hinting varying selective pressures at different loci. The patterns of nucleotide polymorphism are largely determined by the structural and functional constraints of proteins [35]. The gene-wide distributions for pN^(site)^ and pS^(site)^ relative to RSA and DTL highlighted that pN^(site)^ exhibited a distinct inclination for sites with higher RSA and DTL compared to pS^(site)^ (**Fig 7B)**. This result suggests that buried sites (with lower RSA) and functional constraint sites (with lower DTL) are likely to purify nonsynonymous mutation for preserving protein structural stability and function, while revealing comparatively greater tolerance towards synonymous changes. This finding was also supported by the Pearson correlations of log10(pN^(site)^) and log10(pS^(site)^) to RSA and DTL for each gene-sample pair (**Fig. 7B)**, which showed consistently positive correlations for ns-polymorphism sites with an average Pearson coefficient of *r*RSA = 0.22 and *r*DTL = 0.08, whereas the correlations for s-polymorphism sites tended to cluster around 0, averaging *r*RSA = 0.01 and *r*DTL = 0.02. Together, by integrating genetic variants and structural features of RdhA proteins, we found that most polymorphism of *rdhA* genes yielded a strong purifying selection in order to preserve protein stability and metabolic activity of RDases.

## Conclusions

In this study, we present multiple lines of evidence that reductive dehalogenation is a key process supporting the ecology and biogeochemistry of deep-sea cold seeps. Organohalide concentrations reaching up to 18 mg/g, observed dehalogenation activity in laboratory cultures, the widespread presence of both prototypical and transmembrane respiratory reductive dehalogenases, along with the high abundance, expression, and conservation of RDase genes, all suggest that organohalides serve as a central, rather than supplementary, electron acceptor in these environments. Physiologically and phylogenetically diverse bacteria and archaea from some 40 phyla encode RDases, including key players in hydrogen, sulfur, carbon, and nitrogen cycling, with organohalide metabolism likely both directly and indirectly influencing these cycles. It is probable that cold seep microbes have different organohalide substrate preferences, given the chemical diversity of the organohalides available to them and the vast structural diversity of the RDases they encode, which may contribute to metabolic niche partitioning in these competitive environments. In this regard, our analysis also revealed new phylogenetic and structural diversity in the RDases, as well as further insights into the complex evolutionary pathways of these enzymes. Notably, we discovered a novel clade of RDases that contains TMHs, TAT signal peptides, and *rdhB* genes, integrating features of both transmembrane and prototypical respiratory RDases. Finally, our research affirms the deep-sea environment as a repository of novel enzymes, which can be used both directly to remediate organohalide pollutants but also more broadly to advance sequence-structure-function relationships in this space.

## Materials and Methods

### Geochemical and metabolomics analyses

For the analysis of dissolved and solid-phase concentrations of Cl^-^ and Br^-^, samples were measured using a Dionex Ion Chromatograph (Thermo Fisher, USA). The set included 1089 pore water samples and six overlying water samples from 63 sediment cores with lengths of 28, 32, and 40 cm, and 21 longer cores (ranging from 0-320 cm and 0-400 cm) from the Qiongdongnan, Shenhu, Haima, and Site F cold seeps in the South China Sea (**Fig. S1**). Additionally, 68 freeze-dried sediment samples were collected from the Qiongdongnan and Shenhu cold seeps.

For total organic halogens, 55 sediment samples from the Qiongdongnan and Shenhu cold seeps (**Fig. S1**) were analyzed using a multi X® 2500 AOX/TOX analyzer (Analytik Jena, Germany). The untargeted metabolomics analyses of these 55 sediment samples involved processing each sample in a solution of methanol, acetonitrile, and water in a 2:2:1 ratio, with a 20 mg/L internal standard. The samples were vortexed for 30 seconds, ground with steel beads for 10 minutes at 45 Hz, and ultrasonicated for 10 minutes in an ice bath. High-resolution LC-MS/MS analysis of all samples was conducted using a Waters Acquity I-Class PLUS ultra-high performance liquid tandem with a Waters Xevo G2-XS QTOF high-resolution mass spectrometer. Both primary and secondary mass spectrometry data were collected in MSe mode, controlled by the acquisition software (MassLynx V4.2, Waters). The raw data were then processed using Progenesis QI software, based on the METLIN database and Biomark’s self-built library for identification. The relative abundance of metabolites was quantified based on their normalized peak areas, i.e. the percentage of peak area for each metabolite in the total peak area.

### Lab incubations for microbial reductive dehalogenation utilizing various depth sediments along with different halogenated compounds and electron donors

Microcosms were set up using sediment samples from various depths (0-2, 24-26, 124- 126 and 149-151 cmbsf) at the Haima cold seep sites, as previously described [68]. Lactate was used as the electron donor and 2,4,6-tribromophenol (2,4,6-TBP; 25 μM) was added as the electron acceptor. Triplicate abiotic controls for each depth were also conducted using sterilized cold seep sediments. All cultures were incubated in the dark at 30 ± 1°C. Culture media were sampled at different intervals to monitor debromination activity. Samples from the surface sediment microcosms (0-2 cmbsf) were also collected for DNA and RNA extraction. After 25 days of incubation, 10 mL of residual culture medium was collected for DNA extraction and metagenomic sequencing using Illumina NovaSeq platform (Illumina, USA) with PE150 strategy at Personal Co., Ltd. (Shanghai, China). Detailed procedures for DNA and RNA extraction, as well as RT-qPCR for quantifying *rdhA* expression, are provided in the SI Materials and Methods (see Supplementary Materials).

Sediment-free cultures were conducted to test the diversity of dehalogenation substrates and electron donors involved in the cold seep dehalogenation process. These cultures were obtained through ten consecutive transfers of microcosms using surface sediment (5% v/v, 0–2 cmbsf) to a fresh medium amended with 25 μM of 2,4,6-TBP, lactate, and 0.5‰ yeast extract. Perchloroethylene (PCE; 0.65 mM) and 1,1,2-trichloroethane (1,1,2-TCA; 25 mM) were added separately, with headspace samples collected weekly for analysis. In a separate experimental setup, 2,4,6-TBP was used as the substrate, while lactate, acetate, pyruvate, and butyrate were added individually at a concentration of 10 mM to serve as electron donors. Triplicate abiotic controls were conducted for each scenario without microbial inoculation.

For chemical analyses of dehalogenation products from incubation experiments, bromophenols were extracted from the culture media as previously described [69, 70], and analyzed using a gas chromatograph-mass spectrometer (GC-MS) QP2020 (Shimadzu Corporation, Kyoto, Japan) equipped with an SH-Rxi-5Sil MS capillary column (30 m × 0.25 mm inner diameter × 0.25 μm film thickness). Chloroethylenes and chloroethanes were analyzed using an Agilent 7697A headspace autosampler (Agilent Technologies, Santa Clara, USA) connected to an Agilent 8890 GC equipped with a DB-624UI column (30 m × 250 μm × 0.25 μm) and an electron capture detector. Detailed protocols are provided in the SI Materials and Methods.

### Metagenomic and metatranscriptomic datasets from cold seep sediments

Metagenomes were compiled from 165 deep-sea sediment samples, with sediment depths ranging from 0 to 68.55 mbsf and water depths varying from 860 to 3005 meters. These samples were collected from 16 geographically diverse cold seep sites from around the world (**Fig. S1**) [1, 3, 42, 52, 66, 71, 72, 73, 74, 75, 76, 77, 78, 79, 80, 81], encompassing various types of cold seeps such as oil and gas seeps, methane seeps, gas hydrates, asphalt volcanoes, and mud volcanoes. The sites include: Eastern North Pacific (ENP), Santa Monica Mounds (SMM), Western Gulf of Mexico (WGM), Eastern Gulf of Mexico (EGM), Northwestern Gulf of Mexico (NGM), Scotian Basin (SB), Haakon Mosby mud volcano (HM), Mediterranean Sea (MS), Laptev Sea (LS), Jiaolong cold seep (JL), Shenhu area (SH), Haiyang4 (HY4), Qiongdongnan Basin (QDN), Xisha Trough (XST), Haima seep (HM1, HM3, HM5, HM_SQ, S11, SY5, and SY6) and site F cold seep (RS, SF, FR, and SF_SQ). Additionally, 33 metatranscriptomic data were obtained from our previous publications (**Fig. S1**) [52, 66, 79, 82], including those from South China Sea cold seeps in Jiaolong, Haima, Qiongdongnan Basin, and the Shenhu area. The detailed information of the metagenomic and metatranscriptomic datasets used in this study were described in our previous cold seep gene catalog publication [42].

### Metagenomic datasets from other marine sediment environments

Metagenomic samples from various marine sediment environments were retrieved from the Sequence Read Archive database (https://www.ncbi.nlm.nih.gov/sra) by searching for the keyword ‘marine sediment metagenome’ in March 2022. The following criteria were applied for data selection: (1) only paired-end sequencing reads generated by Illumina shotgun platforms were included; (2) datasets involving enriched cultures were excluded; and (3) each dataset had to be larger than 2 Gb. The collected data were then manually categorized based on the isolation source and geographical location, with at least three samples selected from each category. These environments included hadal zone, polar ocean, sinking particle (collected by sediment traps deployed in parallel at 4000 m), and marginal sea. Detailed information on the selected datasets is provided in **Table S22**.

### Metagenomic and metatranscriptomic data processing

The workflow for metagenomic analyses is described in detail (e.g. software and parameters) in **Table S22** [42]. Briefly, metagenomic reads were quality controlled and assembled into contigs. The protein-coding sequences predicted from contigs were clustered at 95% amino acid identity to generate a non-redundant gene catalog (n = 147,289,169). Gene abundances in the non-redundant gene catalog across 178 metagenomes (165 from cold seeps and 13 from other marine sediments) were quantified using Salmon (v.1.9.0) [83] in mapping-based mode (parameters: - validateMappings -meta) and read counts were normalized to GPM (genes per million).

MAGs were derived from assembled contigs of over 1,000 bp using various binning software tools, including MetaBAT2 [84], MaxBin2 [85], CONCOCT [86], SemiBin [87], VAMB [88] and Rosella (https://github.com/rhysnewell/rosella). Produced MAGs were refined, quality controlled, and clustered at 95% average nucleotide identity, resulting in 3,164 species-level representative MAGs. The taxonomy of each MAG was initially assigned using GTDB-TK v2.1.1 with reference to Genome Taxonomy Database R207 [89, 90].

Regarding metatranscriptomes, raw reads were quality filtered (--skip-bmtagger) using Read_QC module within the metaWRAP (v1.3.2) pipeline [91]. To remove ribosomal RNAs from quality-controlled reads, SortMeRNA (v2.1) [92] was used with default settings. Transcript abundances for *rdhA* genes were determined by mapping clean reads from 33 metatranscriptomes to the non-redundant gene catalog using Salmon (v.1.9.0; parameters: -validateMappings -meta) [83]. The transcript abundances of *rdhA* gene were calculated as TPM (transcripts per million).

### Functional annotations and phylogenetic analysis

To identify *rdhA* genes, we used reference *rdhA* sequences (n = 1,040) from the Reductive Dehalogenase Database (https://rdasedb.biozone.utoronto.ca/) [28]. We searched for potential *rdhA* sequences in the non-redundant gene catalog against reference *rdhA* genes in RDaseDB using DIAMOND blastp (v2.0.8) [93], with >30% percentage identity and >50% query coverage as the cut off [43, 53]. Furthermore, we extracted a gene catalog encoding *rdhA* genes from the InterPro database using Pfam id “PF13486” (n = 4,163) and NCBI’s Protein Family Models using NCBI HMM accession “TIGR02486” (n = 1,235). The hmmsearch tool in HMMER v3.3.2 was applied (E-value < 1E-10) using the amino acid sequences of non-redundant gene catalog to the reference gene catalog. To identify potential *rdhA* sequences in MAGs, Prodigal (v2.6.3; parameter: -meta) [94] was utilized to predict protein-coding sequences of all MAGs. These predicted sequences were also searched against the *rdhA* reference sequences using DIAMOND blastp (v2.0.15.153) [93] and HMMER v3.2.1, with the same parameters as mentioned above **(Fig. S8)**. For each gene which passed the required criteria for *rdhA* gene identification by blastp or hmmsearch were merged and then the retrieved results were manually checked according to gene length (> 300 aa) and two Fe-S conserved motifs (CXXCXXCXXXCP, CXXCXXXCP). In brief, pairwise alignment of identified *rdhA* amino acid sequences for conserved active site analysis was performed and visualized using MAFFT v7.471 (-auto option) [95] and Jalview [96]. Phylogenetic trees were further constructed to validate the phylogenetic clades of RdhA, *rdhA* amino acid sequences and reference sequences were aligned using MUSCLE (v3.8.1551, default settings) [97] and trimmed using TrimAL (v1.4.1) [98] with default options. Maximum-likelihood trees were constructed using IQ-TREE (v2.2.0.3) [99] with the “-m MFP -B 1000” options. All trees were visualized by using iTOL (v6) [100].

The MAGs were also annotated using DRAM (v1.3.5; parameter: --min_contig_size 1000) [101] against KEGG, Pfam, MEROPS and dbCAN databases. To annotate genes involved in hydrocarbon degradation, CANT-HYD database [56] with the HMMs of 16 hydrocarbon-degrading genes was searched against with parameters “-cut_nc” using HMMER v3.2.1. The hydrogenases were annotated using DIAMOND blastp (v2.0.15.153; options: --id 50 --query-cover 80 --evalue 1E-20) against local protein databases (https://doi.org/10.26180/c.5230745) [102], further confirmed and classified using the HydDB tool [55]. The VB12Path (https://github.com/qichao1984/VB12Path) was employed to annotate genes involved the VB12 synthesis pathway [59].

### Identification of mobile genetic elements

Classification of *rdhA*-containing contigs as belonging to chromosomes, plasmids or viruses was performed using Genomad v.1.5.0 with default parameters [103]. Integrons, integrative conjugative element (ICEs), IS elements and transposons were identified using HMM searches of the proteins against the 68 marker HMM profiles by default, which are available on proMGE (http://promge.embl.de/) [104].

### Protein modeling and molecular docking

The three-dimensional structures of 586 *rdhA* sequences were generated with AlphaFold (v2.0; full_dbs) [105]. The pLDDT values, a measure for confidence of the AlphaFold structure prediction, ranged from 65 to 97, and averaged at 91. The protein complexes of novel clade RDases were predicted using AlphaFold (v2.0; model_preset = multimer) [105]. The paired alignment of structures relied on 3Di and amino acid based alignment via Foldseek v8.ef4e960 [106]. Structure-based RdhA tree was further built using Foldtree (https://github.com/DessimozLab/fold_tree) based on a local structural alphabet [107] and visualized using iTOL (v6) [100]. Structural similarity and homology relationships of RdhAs with 1,007,623 protein domains in the ECOD database [108] were investigated using Foldseek easy-search module [106] (--tmscore-threshold 0.3 -e 0.001). TMHMM v2.0 was employed to predict transmembrane topology of RdhA proteins (https://services.healthtech.dtu.dk/services/TMHMM-2.0/).

Ledock (v1.0; https://www.lephar.com/) was used to predict the binding poses of 191 halohydrocarbons on different RDases (RMSD: 1.0, Number of binding poses: 20, size: 10). Prediction of ligand channels was performed on the Caver v3.0 (probe_radius 0.9, shell_radius 3, shell_depth 4) [109]. All of the structures were visualized and exported as images using PyMOL (http://www.pymol.org).

### Nucleotide diversity and pN/pS ratio analyses

All metagenomic filtered reads from each sample were mapped to an indexed database of the *rdhA*-containing genomes using Bowtie2 (v2.3.5.1; default parameters) [110]. The nucleotide diversity and pN/pS ratio of *rdhA* genes were calculated from these mappings using the profile module of the inStrain program (v1.6.2; default parameters) [33] at the gene level. To perform gene level profiling, genes were predicted by the software Prodigal (v2.6.3; settings: -p meta) [94] for each MAG carrying *rdhA* genes. A total of 475 *rdhA* genes were retained for microdiversity analyses, which satisfied threshold criteria of having a breadth of 50% and 5× coverage.

### Structure-based polymorphism analyses

The variants across the RdhA protein structures were explored using the microbial population genetics framework implemented in anvi’o (v7.1) [35]. First, population statistics of *rdhA* genes, including coverage, single nucleotide variants (SNVs) and single codon variants (SCVs), were calculated from the mappings using the profile module of the anvi’o program (v7.1; default parameters). Only SNV positions mapping with greater than 10× coverage and the average coverage of *rdhA* genes exceeded 2× in each metagenome were retained (reducing our metagenomic samples size from 165 to 30). Additionally, *rdhA* genes that are present in a minimum of 20 out of 30 metagenomes were included for further analysis. The remaining 200 *rdhA* genes with high-quality structure (pLDDT > 80) were imported for structure-based polymorphism analyses. The SCV data of each site on the high-quality structure (pLDDT > 80) was integrated with ‘anvi-display-structure’, which filtered for variants that had at least 0.05 departure from consensus. The producing results including synonymous (s) and nonsynonymous (ns) polymorphism rates of each codon (pS^(site)^ and pN^(site)^) and RSA. Ligand-binding residues of each gene were predicted with per-residue binding frequencies greater than 0.5 using InteracDome. DTL was calculated using append_dist_to_lig.py (https://merenlab.org/data/anvio-structure/chapter-III/).

### Statistical analyses

Statistical analyses were carried out in R v4.2.3. The normality and variance homogeneity of the data were evaluated using Shapiro-Wilk Test and Levene’s test, respectively. To compare gene abundance and evolutionary metrics among different *rdhA* groups, the Kruskal-Wallis rank-sum test was employed. For paired comparisons within *rdhA* groups, the Wilcoxon test was utilized. Pearson’s product-moment correlation and linear regression were performed to assess the relationship between gene abundance, evolutionary metrics and their relationships with sediment depth. ANOVA analysis was performed to determine the portion of variability in the polymorphism data and the Pearson coefficients for each gene-sample pair were calculated using the R script available at https://merenlab.org/data/anvio-structure/chapter-IV/.

## Data availability

Metagenomic data for the lab-incubated microcosm co-cultured with 2,4,6-TBP has been deposited in the NCBI Sequence Read Archive (SRA) database under the accession number PRJNA1041595. The non-redundant gene and MAGs catalogs derived form 165 cold seep metagenomes can be found in figshare (https://doi.org/10.6084/m9.figshare.22568107). The *rdhA*-carrying MAGs, RdhA protein structures and phylogenetic trees of RdhA based on amino acid sequences and protein structures are available at figshare (https://figshare.com/s/954de2472553e13d69f3).

## Acknowledgements

We thank Fabai Wu, Minghuo Wu, Yinan Deng, Xiaoyan Zhang, Yeting Xie, Qiuyun Jiang, Jing Liao, Chengpeng Li, and Xinyue Liu for helpful discussions and geochemical analyses.

## Author contributions

X.D., Y.H., Z.D., and C.Z. designed this study. X.D., Y.H., Z.D., Y.P., J.P., L.C., D.Z., and Y.X. collected samples, processed the data, performed the geochemical analyses and microbial incubations. Y.H., X.D., Z.D., Y.P., and C.Z. interpreted data. Y.H., Z.D., and Y.P. conducted the data visualization and drafted the manuscript. C.G., C.Z., Y.Y., H.Z., C.Z., D.Z., and M.W. provided useful suggestions to improve the manuscript. X.D. and C.Z. supervised the project. All the authors reviewed the results and approved the manuscript.

## Ethics approval and consent to participate

Not applicable.

